# Sunbeam: an extensible pipeline for analyzing metagenomic sequencing experiments

**DOI:** 10.1101/326363

**Authors:** Erik L. Clarke, Louis J. Taylor, Chunyu Zhao, Andrew Connell, Jung-Jin Lee, Bryton Fett, Frederic D. Bushman, Kyle Bittinger

**Author notes:** These authors contributed equally to this work (Erik L. Clarke, Louis J. Taylor, Chunyu Zhao).

## Abstract

**Background:** Analysis of mixed microbial communities using metagenomic sequencing experiments requires multiple preprocessing and analytical steps to interpret the microbial and genetic composition of samples. Analytical steps include quality control, adapter trimming, host decontamination, metagenomic classification, read assembly, and alignment to reference genomes.

**Results:** We present a modular and user-extensible pipeline called Sunbeam that performs these steps in a consistent and reproducible fashion. It can be installed in a single step, does not require administrative access to the host computer system, and can work with most cluster computing frameworks. We also introduce Komplexity, a software tool to eliminate potentially problematic, low-complexity nucleotide sequences from metagenomic data. Unique components of the Sunbeam pipeline include direct analysis of data from NCBI SRA and an easy-to-use extension framework that enables users to add custom processing or analysis steps directly to the workflow. The pipeline and its extension framework are well documented, in routine use, and regularly updated.

**Conclusions:** Sunbeam provides a foundation to build more in-depth analyses and to enable comparisons in metagenomic sequencing experiments by removing problematic low complexity reads and standardizing post-processing and analytical steps. Sunbeam is written in Python using the Snakemake workflow management software and is freely available at github.com/sunbeam-labs/sunbeam under the GPLv3.

## Background

Metagenomic shotgun sequencing involves isolating DNA from a mixed microbial community of interest, then sequencing deeply into DNAs drawn randomly from the mixture. This is in contrast to marker gene sequencing (e.g., the 16S rRNA gene of bacteria), where a specific target gene region is amplified and sequenced. Metagenomic sequencing has enabled critical insights in microbiology [1–9], especially in the study of virus and bacteriophage communities [10–15]. However, an ongoing challenge is analyzing and interpreting the resulting large datasets in a standard and reliable fashion [16–23].

A common practice to investigate microbial metagenomes is to use Illumina sequencing technology to obtain a large number of short (100-250 base pair) reads from fragmented DNA isolated from a sample of interest. After sequence acquisition, several post-processing steps must be carried out before the sequences can be used to gain insight into the underlying biology [21,24].

Researchers have many tools at their disposal for accomplishing each post-processing step and will frequently encounter multiple parameters in each tool that can change the resulting output and downstream analysis. Varying parameters, tools, or reference database versions between analyses makes it challenging to compare the results of different metagenomic sequencing experiments [25]. Conversely, employing a consistent workflow across studies ensures that experiments are comparable and that the downstream analysis is reproducible, as emphasized in [21]. Documentation of software, databases and parameters used is an essential element of this practice; otherwise, the benefits of consistent and reproducible workflows are lost.

A metagenomic post-processing workflow should have the following qualities to maximize its utility and flexibility: it should be deployable on shared computing systems and in the cloud; it should allow simple configuration of software parameters and reference databases; it should provide error handling and the ability to resume after interruptions; it should be modular so that unnecessary steps can be skipped or ignored, and it should allow new procedures to be added by the user. The ability to deploy the workflow on both institutional and cloud platforms enables workflows to be repeated in different labs with different computing setups and provides flexibility for researchers to choose between computing resources at the institution or in the cloud. Similarly, the ability to record running parameters through the use of configuration files allows for the use of experiment-specific software parameters and serves as documentation for future reference.

Several features contribute to efficient data analysis. It is beneficial if errors or interruptions in the workflow do not require restarting from the beginning: as sequencing experiments produce large amounts of data, having to repeat steps in data processing can be time-consuming and expensive. In addition, not all steps in a workflow will be necessary for all experiments, and some experiments may require custom processing. To handle experiments appropriately, the workflow should provide an easy way to skip unnecessary steps but run them later if necessary. To make the framework widely useful, users must be able to straightforwardly add new steps into the workflow as needed and share them with others. Several pipelines have been developed that achieve many of these goals [26–29], but did not meet our needs for greater flexibility in processing metagenomic datasets and long-term reproducibility of analyses.

Here, we introduce Sunbeam, an easily-deployable and configurable pipeline that produces a consistent set of post-processed files from metagenomic sequencing experiments. Sunbeam is self-contained and installable on GNU/Linux systems without administrator privileges. It features error-handling, task resumption, and parallel computing capabilities thanks to its implementation in the Snakemake workflow language [30]. Nearly all steps are configurable, with reasonable pre-specified defaults, allowing rapid deployment without extensive parameter tuning. Sunbeam can run either using local data directly, or using external data from the National Center for Biotechnology Information (NCBI)’s Sequence Read Archive (SRA) [31]. Sunbeam is extensible using a simple mechanism that allows new procedures to be added by the user.

In addition, Sunbeam features custom software that allows it to process data from challenging sample types, including samples with high proportions of low-quality or host-derived sequences. These include custom-tuned host-derived read removal steps for any number of host or contaminant genomes, and Komplexity, a novel sequence complexity analysis program that rapidly and accurately removes problematic low-complexity reads before downstream analysis. Microsatellite DNA sequences make up a significant proportion of the human genome and are highly variable between individuals [32–34], compounding the difficulty of removing them by alignment against a single reference genome. We developed Komplexity because existing tools for analyzing nucleotide sequence complexity [35–37] did not meet our needs in terms of speed, removal of spurious hits, and natively processing fastq files. We have used Sunbeam in published and ongoing studies of host-associated, low-microbial-biomass body sites [19,38,39], the virome [40], and longitudinal sampling of the microbiome [41,42].

Sunbeam is implemented in Python, Bash, and Rust. It is licensed under the GPLv3. It is freely available at https://github.com/sunbeam-labs/sunbeam and documentation is available at http://sunbeam.readthedocs.io.

## Implementation

### Installation

Sunbeam is installable on GNU/Linux distributions that meet the initial hardware and software requirements listed in Availability and Requirements. Installation is performed by downloading the software from its repository and running “install.sh”. Sunbeam does not require administrator privileges to install or run. We verified that Sunbeam installed and ran a basic analysis workflow on Debian 9; CentOS 6 and 7; Ubuntu 14.04, 16.04, 18.04, and 18.10; Red Hat Enterprise 6 and 7; and SUSE Enterprise 12.

Sunbeam utilizes the Conda package management system [43] to install additional software dependencies beyond those listed in Availability and Requirements. Conda provides automatic dependency resolution, facilitates software installation for non-administrative users, and uses a standardized system for packaging additional software and managing third-party software channels. Conda also provides an isolated environment for additional software dependencies, to avoid conflicts with existing software outside the pipeline. Conda is installed by the Sunbeam installation script if needed. The isolated software environment used by Sunbeam is also created by the installation script.

### Sunbeam architecture

Sunbeam is comprised of a set of discrete steps that take specific files as inputs and produce other files as outputs (Figure 1). Each step is implemented as a rule in the Snakemake workflow framework [30]. A rule specifies the input files needed, the output files produced, and the command or code needed to generate the output files. Implementation of the workflow in Snakemake offered several advantages, including workflow assembly, parallelism, and ability to resume following an interruption. Snakemake determines which steps require outputs from other steps in order to form a directed acyclic graph (DAG) of dependencies when the workflow is started, so the order of execution can change to suit available resources. This dependency DAG prevents ambiguities and cyclic dependencies between steps in the workflow. The DAG structure also allows Snakemake to identify steps that can operate in parallel (e.g. on other processors or compute nodes) if requested by the user. Snakemake manages the scheduling and is compatible with job submission systems on shared clusters.

**Figure 1.**
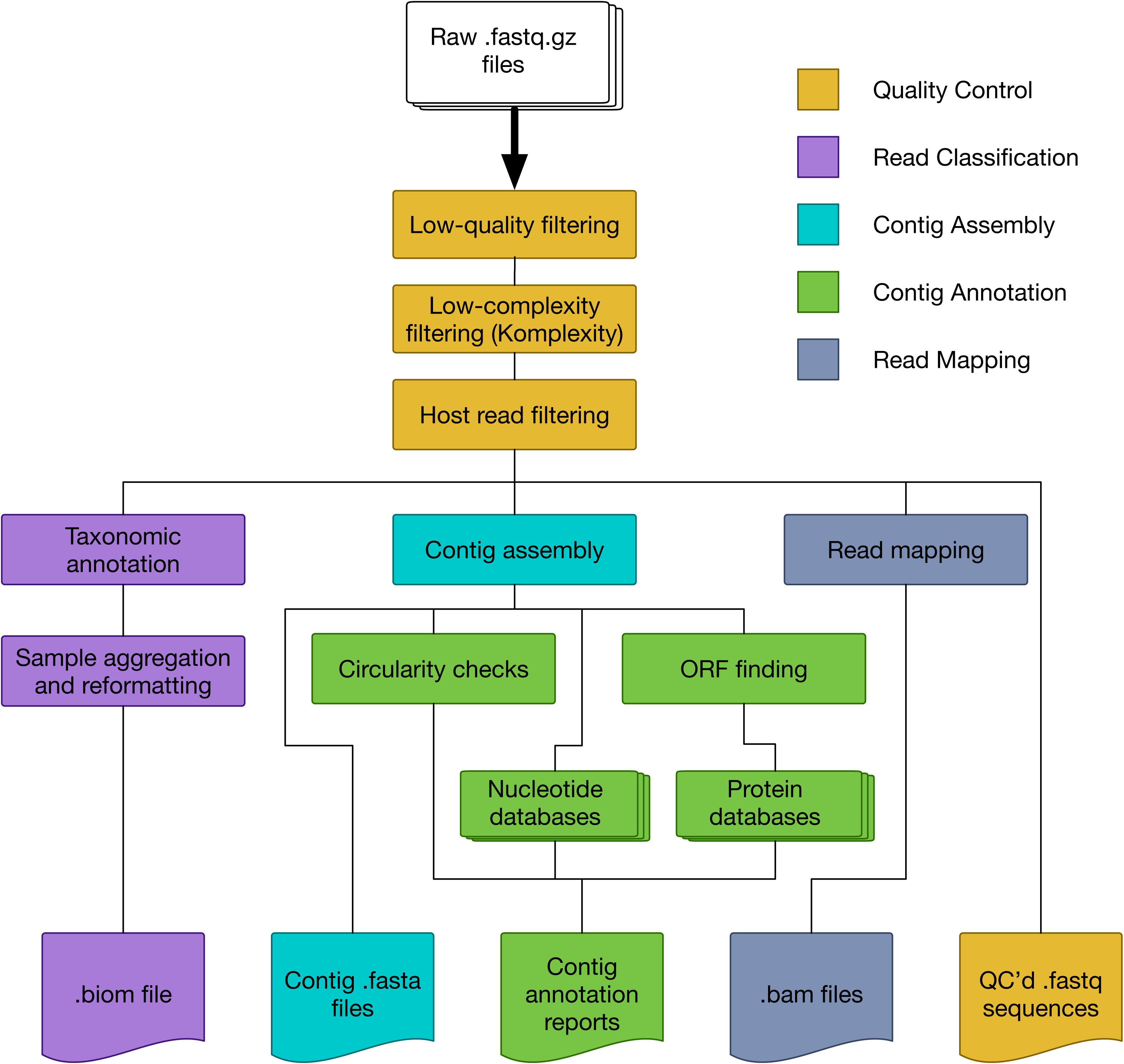
Inputs, processes and outputs for standard steps in the Sunbeam metagenomics pipeline.

The input to the Sunbeam pipeline consists of raw, demultiplexed Illumina sequencing reads, either local files or samples available through the SRA. By default, Sunbeam performs the following preliminary operations on reads in the following order:

1. <u>Quality control: Optionally, reads are downloaded from the SRA using grabseqs</u> [44] <u>and sra-tools</u> [31]. Adapter sequences are removed and bases are quality filtered using the Cutadapt [45] and Trimmomatic [46] software. Read pairs surviving quality filtering are kept. Read quality is assessed using FastQC [47] and summarized in separate reports.
2. <u>Low-complexity masking:</u> Sequence complexity in each read is assessed using Komplexity, a novel complexity scoring algorithm described below. Reads that fall below a user-customizable sequence complexity threshold are removed. Logs of the number of reads removed are written for later inspection.
3. <u>Host read decontamination:</u> Reads are mapped against a user-specified set of host or contaminant sequences using bwa [48]. Reads that map to any of these sequences within certain identity and length thresholds are removed. The numbers of reads removed are logged for later inspection.

After the initial quality-control process, multiple optional downstream steps can be performed in parallel. In the *classify* step, the decontaminated and quality-controlled reads are classified taxonomically using Kraken [49], and summarized in both tab-separated and BIOM [50] format. In the *assembly* step, reads from each sample are assembled into contigs using MEGAHIT [51]. Contigs above a pre-specified length are annotated for circularity. Open reading frames (ORFs) are predicted using Prodigal [52]. The contigs (and associated ORFs) are then searched against any number of user-specified nucleotide or protein BLAST [53] databases, using both the entire contig and the putative ORFs. The results are summarized into reports for each sample. Finally, in the *mapping* step, quality-controlled reads are mapped using bwa [48] to any number of user-specified reference genomes or gene sets, and the resulting BAM files are sorted and indexed using SAMtools [54].

Sunbeam is structured in such a way that the output files are grouped conceptually in different folders, providing a logical output folder structure. Standard outputs from Sunbeam include fastq files from each step of the quality-control process, taxonomic assignments for each read, contigs assembled from each sample, gene predictions, and alignment files of all quality-controlled reads to any number of reference sequences. Most rules produce logs of their operation for later inspection and summary. Users can request specific outputs separately or as a group, and the pipeline will run only the steps required to produce the desired files. This allows the user to skip or re-run any part of the pipeline in a modular fashion.

### Error handling

Most of the actual operations in Sunbeam are executed by third-party bioinformatics software as described above. The error-handling and operational stability of these programs varies widely, and random non-reproducible errors can arise during normal operation. In other cases, execution in a cluster environment can result in resources being temporarily and stochastically unavailable. These circumstances make it essential that Sunbeam can handle these errors in a way that minimizes lost computational effort. This is accomplished in two ways: first, the dependency DAG created by Snakemake allows Sunbeam’s execution to continue with other steps executing in parallel if a step fails. If that step failed due to stochastic reasons, or is interrupted, it can then be re-tried without having to re-execute successful upstream steps. Secondly, Sunbeam wraps execution of many of these programs with safety checks and error-handling code. Where possible, we have identified common failure points and taken measures to allow downstream steps to continue correctly and uninterrupted. For instance, some software will occasionally fail to move the final output file to its expected location. We check for this case and perform the final operation as necessary. In other cases, some programs may break if provided with an empty input file. In rules that use these programs, we have added checks and workarounds to mitigate this problem. Finally, in cases where throwing an error is unavoidable, we halt execution of the rule and provide any error messages generated during the failure. These cases include instances where a step requires more memory than allocated, a variety of error that can be difficult to diagnose. To recover, the user can allocate more memory in the configuration file or on the command line and re-execute.

Sunbeam also inherits all of Snakemake’s error-handling abilities, including the ability to deal with NFS filesystem latency (via the ‘–latency-wait’ option), and automatically re-running failed rules (via the ‘–restart-times’ option). These options are frequently used to overcome issues that arise as part of execution in a cluster environment: in particular, on NFS-based clusters, an upstream rule may complete successfully but the output file(s) are not visible to all nodes, preventing downstream rule execution and an error concerning missing files. The ‘–latency-wait’ option forces Snakemake to wait extra time for the filesystem to catch up. Other workflow systems, such as the Common Workflow Language, allow the workflow to be specified separately from the file paths, and for this reason may offer more safety than the Snakemake framework [55]. In normal use, however, we found that the precautions taken by the Snakemake framework were sufficient to run on shared NFS filesystems at our institutions under typical load.

In addition to runtime error mitigation, Sunbeam performs a series of “pre-flight checks” before performing any operations on the data to preempt errors related to misconfiguration or improper installation. These checks consist of 1) verifying that the environment is correctly configured, 2) that the configuration file version is compatible with the Sunbeam version, and 3) verifying that all file paths specified in the configuration file exist and are accessible.

### Versioning and development

We have incorporated an upgrade and semantic versioning system into Sunbeam. Specifically, the set of output files and configuration file options are treated as fixed between major versions of the pipeline to maintain compatibility. Any changes that would change the format or structure of the output folder, or would break compatibility with previous configuration files, only occur during a major version increase (i.e. from version 1.0.0 to version 2.0.0). Minor changes, optimizations, or bugfixes that do not alter the output structure or configuration file may increase the minor or patch version number (i.e. from v1.0.0 to v1.1.0). Sunbeam stable releases are issued on an as-needed basis when a feature or bug fix is tested and benchmarked sufficiently.

To prevent unexpected errors, the software checks the version of the configuration file before running to ensure compatibility and will stop if it is from a previous major version. To facilitate upgrading between versions of Sunbeam, the same installation script can also install new versions of the pipeline in-place. We provide a utility to upgrade configuration files between major version changes.

To ensure the stability of the output files and expected behavior of the pipeline, we built an integration testing procedure into Sunbeam’s development workflow. This integration test checks that Sunbeam is installable, produces the expected set of output files, and correctly handles various configurations and inputs. The test is run through a continuous integration system that is triggered upon any commit to the Sunbeam software repository, and only changes that pass the integration tests are merged into the ‘stable’ branch used by end-users.

### Extensions

The Sunbeam pipeline can be extended by users to implement new features or to share reproducible reports. Extensions take the form of supplementary rules written in the Snakemake format and define the additional steps to be executed. Optionally, two other files may be provided: one listing additional software requirements, and another giving additional configuration options. Extensions can optionally run in a separate software environment, which enables the use of software dependencies that conflict with Sunbeam’s. Any package available through Conda can be specified as an additional software dependency for the extension. To integrate these extensions, the user copies the files into Sunbeam’s extensions directory, where they are automatically integrated into the workflow during runtime. The extension platform is tested as part of our continuous integration test suite.

User extensions can be as simple or complex as desired and have minimal boilerplate. For example, an extension to run the MetaSPAdes assembly program [56] is shown in Figure 2A. The file *sbx_metaspades_example.rules* specifies the procedure necessary to generate assembly results from a pair of decontaminated, quality controlled FASTQ input files. The pattern for the input files is taken directly from the Sunbeam documentation. The path of the output directory is created by specifying the output directory and the sample name; it was created by modifying the pattern for the standard Sunbeam assembly output, given in the documentation. The shell command at the bottom of the rule is taken directly from the MetaSPAdes documentation at https://biosphere.france-bioinformatique.fr/wikia2/index.php/MetaSPAdes. The extension requires one additional file to run: a file named requirements.txt, containing one line, “spades”, which is the name of the Conda package providing the MetaSPAdes software. In all, the extension comprises a minimal specification of how to install the program, run the program, and name the output files.

**Figure 2.**
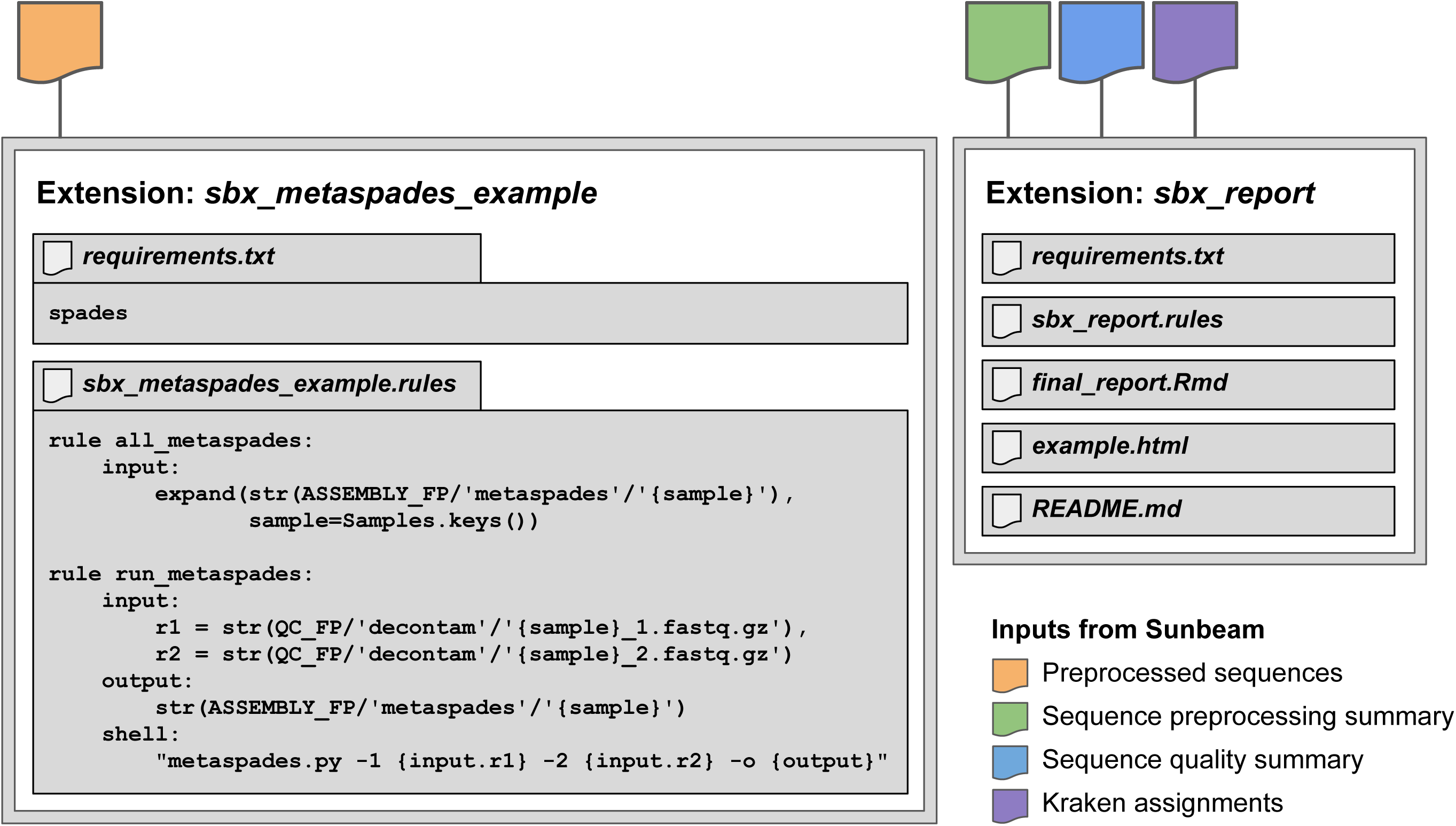
Schematics of example extension inputs and contents. (A) Files for extension *sbx_metaspades_example*, which uses MetaSPAdes to assemble reads from quality-controlled fastq.gz files. *sbx_metaspades_example.rules* lists procedure necessary to generate assembly results from a pair of decontaminated, quality controlled FASTQ input files. *requirements.txt* lists the software requirements for the package to be installed through Conda. (B) Files contained within the *sbx_report* extension: *requirements.txt* lists the software requirements for the package to be installed through Conda; *sbx_report.rules* contains the code for the rule as above, *final_report.Rmd* is a R markdown script that generates and visualizes the report, *example.html* is an example report, and *README.md* provides instructions for installing and running the extension. Sunbeam inputs required for each extension are shown as colored shapes above the extensions.

As a second example, we show the file structure of the *sbx_report* extension, which generates a report from preprocessing, sequence quality, and taxonomic assignment summary files generated in the default workflow (Figure 2B). This extension includes additional files for the report template, an example output report, and a README file with instructions for the user.

Because extensions are integrated directly into the main Sunbeam environment, they have access to the same environmental variables and resources as the primary pipeline, and gain the same error-handling benefits. The manner in which the extensions are integrated into the dependency graph means that a valid extension can only extend, not alter, the primary workflow. Invalid extensions that violate the acyclic dependency requirements will prevent Sunbeam from running. This helps minimize conflicts between extensions as long as unique naming conventions are followed.

To make it easy for users to create their own extensions, we provide documentation, an extension template on our GitHub page (https://github.com/sunbeam-labs/sbx_template), and a number of useful prebuilt extensions available at https://sunbeam-labs.org. We created extensions that allow users to run alternate metagenomic read classifiers like Kaiju [57] or MetaPhlAn2 [58], visualize read mappings to reference genomes with IGV [59], co-assemble contigs from user-specified groups of samples, and even format Sunbeam outputs for use with downstream analysis pipelines like Anvi’o [60]. Users can publish Sunbeam extensions at our website, https://sunbeam-labs.org.

### Komplexity

We regularly encounter low-complexity sequences comprised of short nucleotide repeats that pose problems for downstream taxonomic assignment and assembly [12,39], for example by generating spurious alignments to unrelated repeated sequences in database genomes. These are especially common in low-microbial biomass samples associated with vertebrate hosts. To avoid these potential artifacts, we created a novel, fast read filter called Komplexity. Komplexity is an independent program, implemented in the Rust programming language for rapid, standalone performance, designed to mask or remove problematic low-complexity nucleotide sequences. It scores sequence complexity by calculating the number of unique k-mers divided by the sequence length. Example complexity score distributions for reads from ten stool virome samples (high microbial biomass; [15]) and ten bronchoalveolar lavage (BAL) virome samples (low-biomass, high-host; [12]) are shown in Figure 4B—low-complexity reads are often especially problematic in low-microbial-biomass samples like BAL. Komplexity can either return this complexity score for the entire sequence or mask regions that fall below a score threshold. The k-mer length, window length, and complexity score cutoff are modifiable by the user, though default values are provided (k-mer length = 4, window length = 32, threshold = 0.55). Komplexity accepts FASTA and FASTQ files as input and outputs either complexity scores or masked sequences in the input format. As integrated in the Sunbeam workflow, Komplexity assesses the total read complexity and removes reads that fall below the default threshold. Although low-complexity reads are filtered by default, users can turn off this filtering or modify the threshold in the Sunbeam configuration file. Komplexity is also available as a separate open-source program at https://github.com/eclarke/komplexity.

## Results and Discussion

Sunbeam implements a core set of commonly-required tasks supplemented by user-built extensions. Even so, the capabilities of Sunbeam compare favorably with features offered in existing pipelines such as SURPI (Sequence-based Ultra-Rapid Pathogen Identification) [26], EDGE (Empowering the Development of Genomics Expertise) [27], ATLAS (Automatic Tool for Local Assembly Structures) [28], and KneadData [29]. A detailed feature comparison is shown in Table 1. Where Sunbeam’s primary advancements lie are in its extension framework and novel software to address the issues of low-complexity or host-derived sequence filtering. The Sunbeam extensions framework facilitates the addition of new features to Sunbeam without adding overhead to the core pipeline—once developed, a Sunbeam extension can be discovered and used by anyone through our extensions website (www.sunbeam-labs.org). Sunbeam extensions that generate figures can also be used to promote reproducible science (examples below). As Sunbeam can work directly from SRA data, regenerating figures and analyses from a study is as painless as installing the extension, initializing using the BioProject or SRA project identifier, then running the pipeline as usual.

**Table 1.** – Feature comparison for metagenomic pipelines.

|  | **Sunbeam** | **SURPI** | **KneadData** | **EDGE** | **ATLAS** |
| --- | --- | --- | --- | --- | --- |
| **Architecture/usage** |  |  |  |  |  |
| Dependency management | Conda | Bash | Pip (partial) |  | Conda |
| Modularity | Snakemake |  |  | Perl modules | Snakemake |
| Results reporting | Tables, coverage maps, figures | Tables, coverage maps |  |  | Tables, coverage maps |
| Extension framework | Sunbeam extensions |  |  |  |  |
| Clinical certification |  | CLIA |  |  |  |
| Data source | Local, SRA | Local | Local | Local | Local |
| **Quality control** |  |  |  |  |  |
| Adapter trimming | Trimmomatic, Cutadapt | Cutadapt | Trimmomatic | FaQCs | BBDuk2 |
| Error correction |  |  |  |  | Tadpole |
| Read quality | Fastqc | Cutadapt | Fastqc | FaQCs | BBDuk2 |
| Host filtering | Any | Human | Any | Any | Any |
| Low-complexity | Komplexity | DUST | TRF | Mono- or dinucleotide repeats | BBDuk2 |
| Read subsampling/rarefaction | VSEARCH (extension) |  |  |  |  |
| **Sequence analysis** |  |  |  |  |  |
| Reference alignment | BWA |  |  | Bowtie2, MUMmer+ JBrowse | BBMap |
| Classification | Kraken, (MetaPhlAn2, Kaiju extensions) | SNAP |  | GOTTCHA, Kraken, MetaPhlAn | DIAMOND |
| Assembly | MEGAHIT | Minimo |  | IDBA-UD, SPAdes | MEGAHIT, SPAdes |
| ORFs (aa) | Prodigal, BLASTp |  |  |  | Prokka |
| Full contig (nt) | Circularity, BLASTn | RAPSearch |  | BWA | DIAMOND |
| Functional annotation | eggNOG (extension) |  |  |  | ENZYME/eggNOG/dbCAN |
| Phylogeny reconstruction |  |  |  | PhaMe, FastTree/RAXML |  |
| Primer design |  |  |  | BW, Primer3 |  |

Feature comparison for metagenomic pipelines. Tools used by each pipeline: trimmomatic [46]; cutadapt [45];
tadpole [61]; fastqc [47]; FaQCs [62]; BBDuk2 [63]; DUST [36]; TRF [64]; VSEARCH [65]; bwa [48]; bowtie2 [66];
BBMap [67]; KRAKEN [49]; SNAP [68]; MUMmer [69]; JBrowse [70]; GOTTCHA [71]; MetaPhlAn [58]; DIAMOND
[72]; FastTree [73]; MEGAHIT [51]; SPAdes [74]; Minimo [75]; Prodigal [52]; BLASTp [53]; Prokka [76]; BLASTn [53]; eggNOG [77]; ENZYME [78]; dbCAN [79]; Primer3 [80]; RAPSearch [81]; RAxML [82]; conda [43]; PhaME [83]; Snakemake [30]; SAMtools [54].

To demonstrate the development of extensions and the use of Sunbeam on real-world data, we reproduced key results from three published studies and tested Sunbeam on an internal pilot study of shallow shotgun sequencing (Figure 3). These studies span multiple research areas (high-biomass human gut microbiome, soil microbiome, virome, and shallow shotgun methodology) and emphasize Sunbeam’s versatility. Each of these analyses is packaged and distributed as a Sunbeam extension. The extension workflow begins with downloading data from the SRA and ends with generation of a final report. We verified that the example workflows produced identical results on our institutional systems and on cloud-based systems.

**Figure 3.**
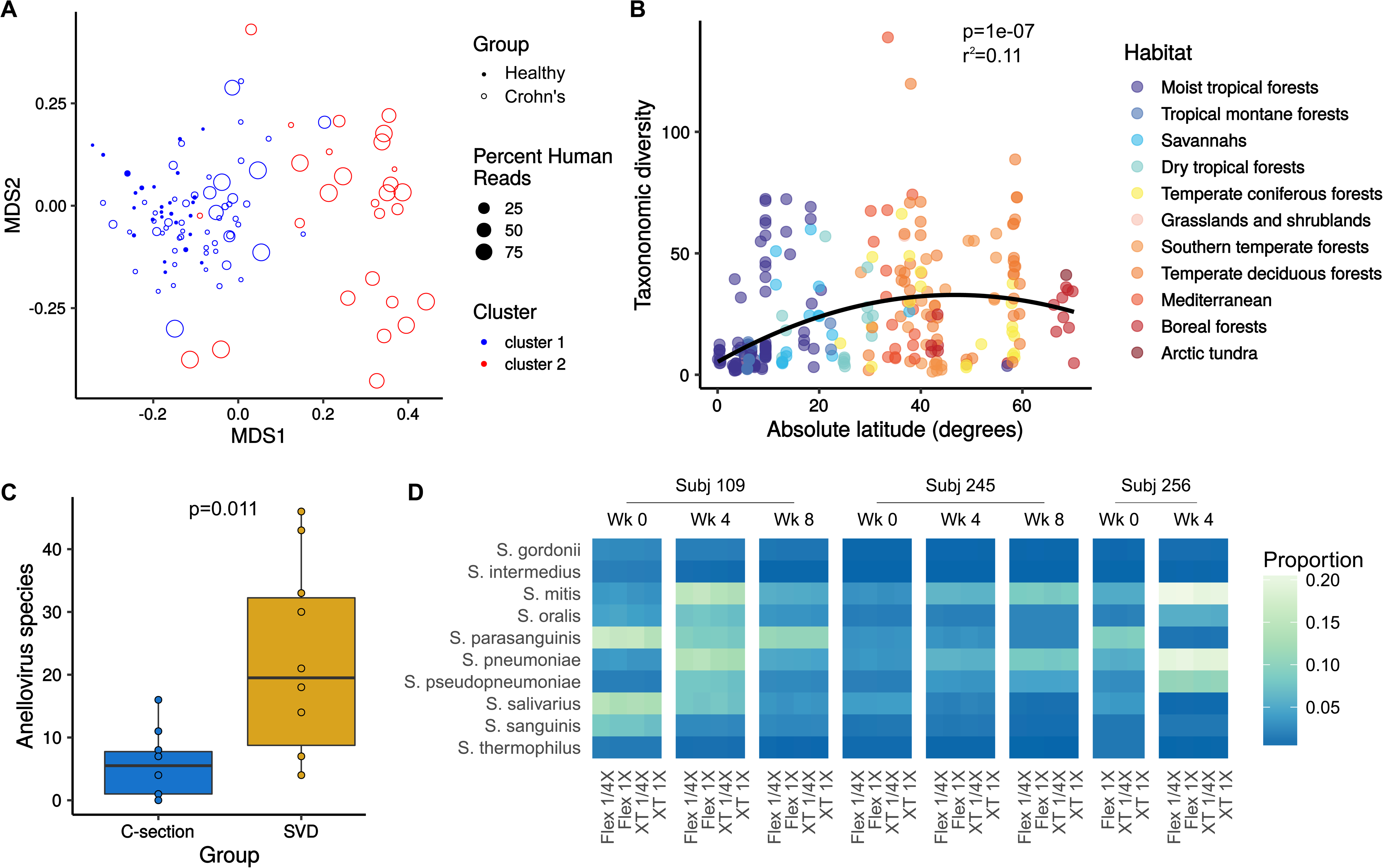
(A) Nonmetric Multidimensional Scaling plots generated using the vegan package in R [99], using MetaPhlAn2 classifications of data from Lewis *et al*. 2015 [84]. Each point is colored by the cluster in which it was annotated in the Lewis *et al*. metadata—cluster 2 (red) is the dysbiotic cluster, while cluster 1 (blue) is the healthy-like cluster. (B) Inverse Simpson diversity by absolute latitude calculated using the vegan package in R from the Kraken classification output of Sunbeam for Bahram *et al*. 2018 [86]. Points are colored by habitat. The polynomial regression line is shown in black. (C) Boxplots of unique *Anelloviridae* taxa in each sample from McCann *et al*. 2018 [87]. Each point corresponds to a single sample. (D) Heatmap from shallow shotgun analysis colored by proportional abundance. Each row corresponds to a bacterial taxon; each column represents a different reagent combination. Columns are grouped by timepoint, then by subject (top). All plots were generated using the ggplot2 R package [100].

In Figure 3A, we reproduce a finding from Lewis *et al*. 2015 showing distinct gut microbial community clusters in individuals with Crohn’s disease, which track with human DNA abundance in stool [84]. As this finding depends on the use of read-based classification, we used Kraken (the Sunbeam built-in classifier) as well as extensions for Kaiju [57] and MetaPhlAn2 [58] to test the consistency of the conclusions across different tools—the original study used MetaPhlAn [85] to classify reads. All three classification methods recapitulated the previously reported clusters (MetaPhlAn2 results in Figure 3A, full report in Additional File 1). This analysis can be re-run using the sbx_lewis2015 extension (https://github.com/louiejtaylor/sbx_lewis2015).

An example of a non-host-associated analysis is shown in Figure 3B. Bahram *et al*. 2018 showed that soil bacterial diversity was highest in temperate latitudes, but lower at the equator and in artic regions [86]. We used Sunbeam to process and classify reads from this dataset and found similar results (Figure 2B; *P*<0.001, *R*^2^=0.11; Bahram *et al*. *P*<0.001, *R*^2^=0.16). This analysis can be reproduced using the sbx_bahram2018 extension (https://github.com/louiejtaylor/sbx_bahram2018, Additional File 2).

In Figure 3C, we reproduce results from the virome study published by McCann et al. 2018, which found differences according to mode of birth in gut viral communities of one-year-olds [87]. One salient finding was that *Anelloviridae*, a family of ubiquitous human commensal viruses [88], were much more diverse in children born by spontaneous vaginal delivery (SVD) compared to those born via C-section. We used Sunbeam to classify the reads from this dataset and also found more unique anelloviruses in SVD compared to C-section (Figure 3C, Additional File 3; *P*=0.011; McCann *et al*. *P*=0.014). The finding of McCann *et al* was recovered using different approaches to identify viruses: McCann *et al*. used a translated nucleotide query against a database of anellovirus ORF1 proteins, while we used Kraken-based classification. This analysis can be reproduced using the sbx_mccann2018 extension (https://github.com/louiejtaylor/sbx_mccann2018).

Figure 3D shows results from a methods development pilot study conducted at the CHOP Microbiome Center. Here, we sequenced tongue swab specimens from three healthy children, collected at four-week intervals. DNA from the tongue swabs was prepared for sequencing using two library preparation kits: the Nextera XT kit and the Nextera DNA Flex kit. For each kit and specimen, the DNA library was prepared using a full-scale (1X) and quarter-scale (1/4X) reagent volumes relative to standard Illumina protocols. We conducted shotgun metagenomic sequencing and recovered approximately 500,000 reads per sample, after host filtering. Thus, our pilot study is an example of “shallow shotgun sequencing,” which is emerging as a cost-effective alternative to 16S rRNA marker gene studies [89]. In our analysis of the pilot study, the type of kit had a small but measurable effect on the abundance of Streptococcus species, but the estimated effect size for the kit (*R*^2^ = 0.003 for *S. mitis*) was orders of magnitude less than that between specimens (*R*^2^ = 0.995). This analysis can be reproduced using the sbx_shallowshotgun_pilot extension (https://github.com/junglee0713/sbx_shallowshotgun_pilot; report HTML file in Supplementary File 4).

Sunbeam’s extension framework promotes reproducible analyses and greatly simplifies performing the same type of analysis on multiple datasets. Extension templates, as well as a number of pre-built extensions for metagenomic analysis and visualization software like Anvi’o [60], MetaPhlAn [58], and Pavian [90], are available on our GitHub page (https://github.com/sunbeam-labs). Sunbeam’s ability to use data directly from the SRA facilitates reproducibility: final figure generation can be integrated into a Sunbeam extension, greatly lowering the barrier to reproducing analyses and studies.

### Comparing low-complexity filtering program filtering and performance

Low complexity reads often cross-align between genomes, and commonly elude standard filtersin use today. The gold standard of such filtering programs, RepeatMasker [35], uses multiple approaches to identify and mask repeat or low complexity DNA sequences, including querying a database of repetitive DNA elements (either Repbase [91] or Dfam [92]). Another program, used in the BLAST+ suite, DUST [36] employs an algorithm which scores and masks nucleotide sequence windows that exceed a particular complexity score threshold (with lower-complexity sequences assigned higher scores) such that no subsequence within the masked region has a higher complexity score than the entire masked region. BBMask, developed by the Joint Genome Institute, masks sequences that fall below a threshold of k-mer Shannon diversity [37].

Many of these tools were not optimal for our use with shotgun metagenomic datasets. RepeatMasker uses databases of known repeat sequences to mask repetitive nucleotide sequences, but runs too slowly to be feasible for processing large datasets. An option to filter reads falling below a certain complexity threshold is not available in DUST, RepeatMasker or BBMask (although filtering is available in the BBMask companion tool BBDuk). Finally, the memory footprint of BBMask scales with dataset size, requiring considerable resources to process large shotgun sequencing studies. Therefore, we designed Komplexity to mask or filter metagenomic reads as a rapid, scalable addition to the Sunbeam workflow that can also be installed and run separately. It accepts FASTA/Q files as input, can mask or remove reads below a specified threshold, and operates with a constant memory footprint. Our goal was to achieve quality comparable to RepeatMasker in a reasonable timeframe.

To compare the performance of all the low-complexity-filtering tools discussed above, we used pIRS [93] to simulate Illumina reads from the human conserved coding sequence dataset [94] as well as human microsatellite records from the NCBI nucleotide database [95] with the following parameters: average insert length of 170 nucleotides with a 5% standard deviation, read length of 100 nucleotides, and 5x coverage. To ensure compatibility with all programs, we converted the resulting files to FASTA format, then selected equal numbers of reads from both datasets for a total of approximately 1.1 million bases in the simulated dataset (available at https://zenodo.org/record/2541222) [96]. We processed the reads using Komplexity, RepeatMasker, DUST and BBMask and used GNU Time version 1.8 [97] to measure peak memory usage and execution time for six replicates (Table 2). Komplexity and RepeatMasker mask a similar proportion of microsatellite nucleotides, while none of the four tools masks a large proportion of coding nucleotides. Komplexity runs faster and has a smaller memory footprint than other low-complexity filtering programs. The memory footprint of Komplexity and DUST are also relatively constant across datasets of different sizes (data not shown).

**Table 2.**
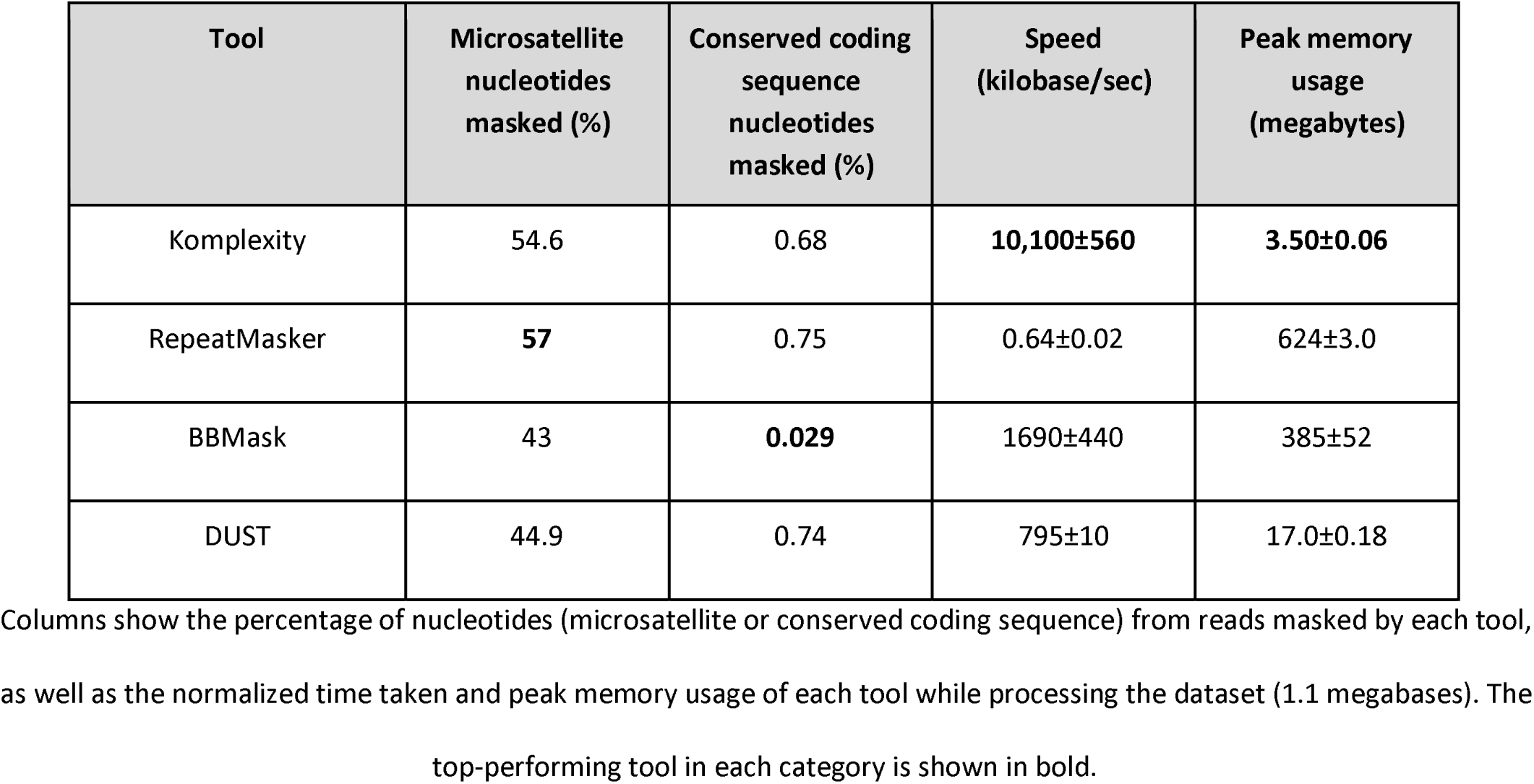
Memory usage, speed, and nucleotides masked for each program.

| Tool | Microsatellite nucleotides masked (%) | Conserved coding sequence nucleotides masked (%) | Speed (kilobase/sec) | Peak memory usage (megabytes) |
| --- | --- | --- | --- | --- |
| Komplexity | 54.6 | 0.68 | **10,100±560** | **3.50±0.06** |
| RepeatMasker | **57** | 0.75 | 0.64±0.02 | 624±3.0 |
| BBMask | 43 | **0.029** | 1690±440 | 385±52 |
| DUST | 44.9 | 0.74 | 795±10 | 17.0±0.18 |

Columns show the percentage of nucleotides (microsatellite or conserved coding sequence) from reads masked by each tool, as well as the normalized time taken and peak memory usage of each tool while processing the dataset (1.1 megabases). The top-performing tool in each category is shown in bold.

To understand the extent to which different tools might synergize to mask a larger proportion of overall nucleotides, we visualized nucleotides from the microsatellite dataset masked by each tool or combinations of multiple tools using UpSetR [98] (Figure 4A). Komplexity masks 78% of the nucleotides masked by any tool, and 96% excluding nucleotides masked by only RepeatMasker. This suggests that there would only be a marginal benefit to running other tools in series with Komplexity. Komplexity in combination with Sunbeam’s standard host removal system resulted in the removal of over 99% of the total simulated microsatellite reads.

**Figure 4.**
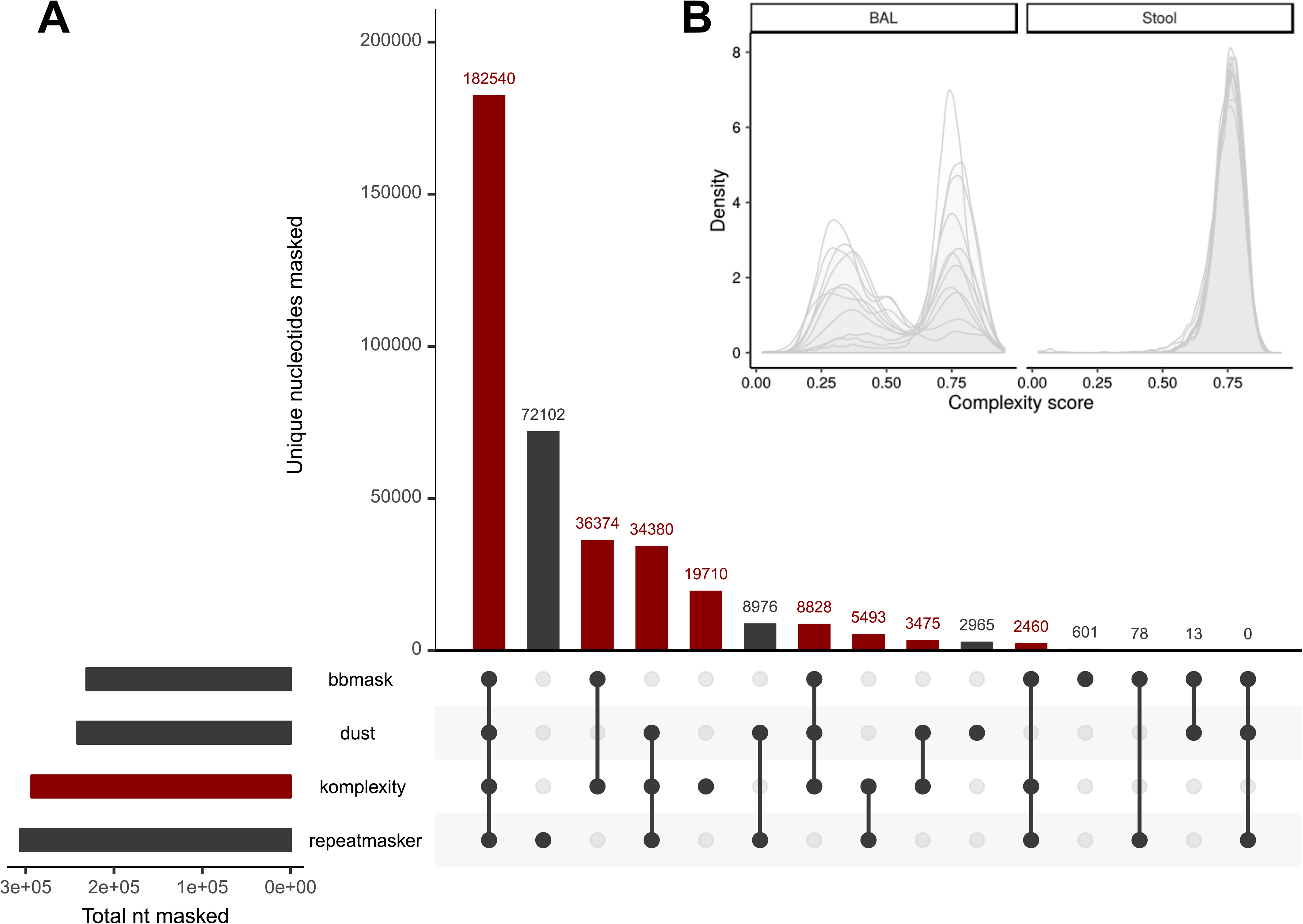
(A) Comparison between Komplexity and similar software (BBMask, DUST, and RepeatMasker). The small bar plot in the lower left shows the total nucleotides masked by each tool. The central bar plot shows the number of unique nucleotides masked by every tool combination; each combination is shown by the connected dots below. Bars displaying nucleotides masked by tool combinations that include Komplexity are colored red. (B) Example complexity score distributions calculated by Komplexity for reads from ten stool virome samples (high microbial biomass; [15]) and ten bronchoalveolar lavage (BAL) virome samples (low-biomass, high-host; [12]) using the default parameters.

## Conclusions

Here we introduce Sunbeam, a Snakemake-based pipeline for analyzing shotgun metagenomic data with a focus on reproducible analysis, ease of deployment and use. We also present Komplexity, a tool for rapidly filtering and masking low-complexity sequences from metagenomic sequence data, and show its superior performance in comparison with other tools for masking human microsatellite repeat sequences. Sunbeam’s scalability, customizability, and facilities for deployment simplify the processing of shotgun metagenomic sequence data, while its extension framework enables customized reproducible analyses. We have already used Sunbeam in multiple published [19,38–42] and ongoing studies.

## Availability and requirements

**Project name:** Sunbeam

**Project home page:** https://github.com/sunbeam-labs/sunbeam

**Operating system:** GNU/Linux; verified on the following distributions: Debian 9; CentOS 6 and 7; Ubuntu 14.04, 16.04, 18.04, and 18.10; Red Hat Enterprise 6 and 7

**Programming languages:** Python, Rust, and Bash

**Other requirements:** Software: Python (version 2.7, 3.4, 3.5, or 3.6), git (version >= 2), GNU Coreutils, wget, bzip2, and Bash (version >= 3) required for installation. At least 100 GB hard drive space and 16 GB memory are recommended to run the pipeline, dependent on databases and input file sizes used.

**License:** GPLv3

**Restrictions to use by non-academics:** No

### List of abbreviations

ATLAS: Automatic Tool for Local Assembly Structures
BAM: binary alignment map
BLAST: Basic Local Alignment Search Tool
DAG: Directed Acyclic Graph
EDGE: Empowering the Development of Genomics Expertise
GPL: (GNU) General Public License
ORF(s): open reading frame(s)
SDUST: Symmetric DUST
SRA: Sequence Read Archive
SURPI: Sequence-based Ultra-Rapid Pathogen Identification
SVD: Spontaneous Vaginal Delivery

## Declarations

### Ethics approval and consent to participate

The study protocol was approved by The Children's Hospital of Philadelphia Institutional Review Board. All subjects gave written, informed consent prior to participation.

### Consent for publication

Not applicable.

### Availability of data and material

Sunbeam is available at https://github.com/sunbeam-labs/sunbeam. Komplexity is available at https://github.com/eclarke/komplexity. Pre-built extensions referenced can be found at https://github.com/sunbeam-labs/. The dataset of simulated microsatellite and conserved coding sequence reads is archived in Zenodo at https://zenodo.org/record/2541222. Shotgun data from the shallow shotgun pilot study is deposited in SRA under project identifier SRP178101. The analyses in Figure 3 can be reproduced by the following Sunbeam extensions: https://github.com/louiejtaylor/sbx_lewis2015, https://github.com/louiejtaylor/sbx_bahram2018, https://github.com/louiejtaylor/sbx_mccann2018, https://github.com/junglee0713/sbx_shallowshotgun_pilot.

### Competing interests

The authors declare that they have no competing interests.

### Funding

This work was supported by the NIH grants U01HL112712 (Site-Specific Genomic Research in Alpha-1 Antitrypsin Deficiency and Sarcoidosis (GRADS) Study), R01HL113252, and R61HL137063, and received assistance from the Penn Center for AIDS Research (P30AI045008), T32 Training Grant (T32AI007324, LJT), and the PennCHOP Microbiome Program (Tobacco Formula grant under the Commonwealth Universal Research Enhancement (C.U.R.E) program with the grant number SAP # 4100068710).

### Authors’ contributions

ELC, CZ, FDB, and KB conceived and designed Sunbeam. ELC, LJT, CZ, AC, and KB developed Sunbeam. ELC and LJT conceived and developed Komplexity. LJT performed the low-complexity sequence masking analysis, reproduced results from previously published studies, and wrote the associated extensions. ELC, LJT, AC, and KB wrote the manuscript. BF conducted the shallow shotgun pilot study presented here; JJL conducted the analysis and wrote the shallow shotgun Sunbeam extension. All authors read, improved, and approved the final manuscript.

## Acknowledgements

Thanks to members of the Bushman lab, Penn-CHOP Microbiome Center, and Penn Bioinformatics Code Review communities for helpful suggestions, discussions, and beta testing.

## Additional file legends

### Additional File 1 (HTML): Lewis *et al*. 2015 Report

Report with figures reproducing results from Lewis *et al*. 2015 [84].

### Additional File 2 (HTML): Bahram *et al*. 2018 Report

Report with figures reproducing results from Bahram *et al*. 2018 [86].

### Additional File 3 (HTML): McCann *et al*. 2018 Report

Report with figures reproducing results from McCann *et al*. 2018 [87].

### Additional File 4 (HTML): Shallow Shotgun Pilot Report

Report with figures relating to the shallow shotgun pilot study described above.

