## Additional File 1 for "Sunbeam: an extensible pipeline for analyzing metagenomic sequencing experiments"

Lewis et al 2015: Crohn�s disease clusters


### Lewis *et al* 2015: Crohn�s disease clusters

###### *10 January, 2019*

#### Background

This report uses the Sunbeam pipeline to reproduce findings from the 2015 Lewis *et al.* paper entitled �Inflammation, Antibiotics, and Diet as Environmental Stressors of the Gut Microbiome in Pediatric Crohn�s Disease� (PMID: 26468751). A key finding of this paper was that individuals with Crohn�s disease form two distinct clusters by Multidimensional Scaling (MDS)�the dysbiotic cluster tended to have a high human DNA fraction. Here, we test whether we can reproduce this finding using the originial data submitted to the SRA, and compare three different read-based classification methods available in Sunbeam or as Sunbeam extensions: Kaiju, Kraken, and MetaPhlAn2.

This report was generated by the extension sbx\_lewis2015; this link also includes instructions for re-running this analysis from the beginning. It depends on output from two other extensions: sbx\_kaiju and sbx\_metaphlan.

#### Results

Below are Nonmetric Multidimensional Scaling plots generated using the `vegan` package in R. Each point is colored by the cluster in which it was annotated in the Lewis *et al* metadata, to query whether our results match those published previously. Cluster 2 (red) is the dysbiotic cluster, while cluster 1 (blue) is the healthy-like cluster. Lewis *et al* used MetaPhlAn to classify reads in the original analysis.

##### Kraken results

##### Kaiju results

##### MetaPhlAn2 results

For these, a few samples came out of MetaPhlAn with 100% unclassified�they are omitted from this plot.

#### Conclusion

All three classification methods support the conclusions of the original paper.
