## Additional File 2 for "Sunbeam: an extensible pipeline for analyzing metagenomic sequencing experiments"

Bahram et al 2018: Soil bacterial diversity


### Bahram *et al* 2018: Soil bacterial diversity

###### *10 January, 2019*

#### Background

This report uses the Sunbeam pipeline to reproduce findings from �Structure and function of the global topsoil microbiome� (PMID: 30069051) by Bahram *et al*. A key finding of this paper is that bacterial diversity is highest in temperate habitats, but lower in extreme latitudes and near the equator. This extension tests whether we can reproduce this finding using Sunbeam and Kraken.

This report was generated by the extension sbx\_bahram2018; this link also includes instructions for re-running this analysis from the beginning.

#### Results

Using the Kraken output of Sunbeam (rarefied as in Bahram *et al*), we plot inverse Simpson diversity by absolute latitude calculated using the `vegan` package in R. Points are colored by habitat.

##### Kraken results

#### Conclusion

The results of our analysis are comparable to those reported by Bahram *et al*. The polynomial regression p-value (p=1.068456e-07) and adjusted r-squared (r^2^=0.1068863) for the relationship between inverse Simpson diversity and absolute latitude are comparable to the values reported in the manuscript (p=1e-07; r^2^=.160).
