## Additional File 3 for "Sunbeam: an extensible pipeline for analyzing metagenomic sequencing experiments"

McCann et al 2018: Post-FMT viral taxa


### McCann *et al* 2018: Post-FMT viral taxa

###### *13 January, 2019*

#### Background

This report uses the Sunbeam pipeline to reproduce findings from �Viromes of one year old infants reveal the impact of birth mode on microbiome diversity� (PMID: 29761040) by Zuo *et al*. A key finding of this study is that the virome at 1 y/o seems to correlate with the birth mode�that is, that children born by spontaneous vaginal delivery (SVD) have higher viral diversity and higher *Anelloviridae* richness than children born by C-section. The purpose of this report is to see whether analysis with Sunbeam reproduces these results. This report was generated by the extension sbx\_mccann2018; this link also includes instructions for re-running this analysis from the beginning. This report was run using the Kraken Standard database built 1 October, 2018.

#### Results

Below are boxplots generated using the `ggplot2` package in R. Each point corresponds to a single sample. We want to see whether we also find greater Anellovirus richness in spontaneous vaginal delivery (SVD) compared to C-section, and greater overall virus diversity in SVD compared to C-section:

##### Anellovirus richness

##### Virus diversity

#### Conclusion

The results of our analysis are comparable to those reported by McCann *et al*. The Wilcoxon rank-sum test p-value (p=0.01102403) is similar to that reported in the paper (p=0.014). While the difference in viral diversity does not reach significance (p=0.1035892)), we do see the same relationship in Shannon diversity (higher diversity in SVD) as McCann *et al*.
