## Additional File 4 for "Sunbeam: an extensible pipeline for analyzing metagenomic sequencing experiments"

CHOP Shallow Shotgun Pilot Study


### CHOP Shallow Shotgun Pilot Study

Tongue swab samples were collected from three healthy subjects at baseline, four weeks, and eight weeks from study initiation. DNA was extracted with the Qiagen DNeasy PowerSoil kit, and was prepared for shotgun metagenomic sequencing using either the Nextera XT kit (XT) or Nextera Flex DNA kit (Flex). Each kit was used with both the full reagent volume (1X), or with 1/4 reagent volume (1/4X). Libraries were sequenced on an Illumina HiSeq 2500 instrument.

### Average nucleotide quality

Average nucleotide quality score after trimming adapter and filtering low quality (only forward read or reverse read or neither was kept after trimmomatic) read pairs is given. Red dots represent the mean quality score and vertical line segments represent mean \(\pm\) sample standard deviation.

### Read Counts

The dashed line represents 1 million read counts.

### Heatmap of taxonomic assignments

Here, we focus on Streptococcus, the predominant genus.

```
## 
## Call:
## lm(formula = log10(Proportion) ~ Specimen + lib_prep + rxn_scale, 
##     data = .)
## 
## Residuals:
##       Min        1Q    Median        3Q       Max 
## -0.030765 -0.007870  0.001439  0.006046  0.025389 
## 
## Coefficients:
##                        Estimate Std. Error  t value Pr(>|t|)    
## (Intercept)           -0.820360   0.008081 -101.517  < 2e-16 ***
## SpecimenSpecimen 1064 -0.442511   0.010130  -43.683  < 2e-16 ***
## SpecimenSpecimen 1897 -0.654976   0.010130  -64.657  < 2e-16 ***
## SpecimenSpecimen 2005 -0.444399   0.011013  -40.354  < 2e-16 ***
## SpecimenSpecimen 2132 -0.238782   0.010130  -23.572  < 2e-16 ***
## SpecimenSpecimen 2134 -0.377983   0.010130  -37.313  < 2e-16 ***
## SpecimenSpecimen 2151  0.142903   0.010130   14.107 3.49e-12 ***
## SpecimenSpecimen 820  -0.620535   0.010130  -61.257  < 2e-16 ***
## lib_prepXT            -0.027622   0.005179   -5.334 2.74e-05 ***
## rxn_scale1X            0.001416   0.005179    0.273    0.787    
## ---
## Signif. codes:  0 '***' 0.001 '**' 0.01 '*' 0.05 '.' 0.1 ' ' 1
## 
## Residual standard error: 0.01433 on 21 degrees of freedom
## Multiple R-squared:  0.9981, Adjusted R-squared:  0.9972 
## F-statistic:  1208 on 9 and 21 DF,  p-value: < 2.2e-16
```

```
## Analysis of Variance Table
## 
## Response: log10(Proportion)
##           Df  Sum Sq Mean Sq   F value  Pr(>F) R-squared
## Specimen   7 2.22524 0.31789 1548.9013 0.00000   0.99544
## lib_prep   1 0.00588 0.00588   28.6275 0.00003   0.00263
## rxn_scale  1 0.00002 0.00002    0.0747 0.78724   0.00001
## Residuals 21 0.00431 0.00021                     0.00193
```

### Alpha diversity

Alpha diversity (within sample diversity) was assessd by the Shannon index.

```
## 
## Call:
## lm(formula = ShannonIdx ~ Specimen + lib_prep + rxn_scale, data = alpha_df)
## 
## Residuals:
##      Min       1Q   Median       3Q      Max 
## -0.12329 -0.05014 -0.01370  0.03130  0.21555 
## 
## Coefficients:
##                        Estimate Std. Error t value Pr(>|t|)    
## (Intercept)            3.101521   0.050256  61.714  < 2e-16 ***
## SpecimenSpecimen 1064 -0.202864   0.063000  -3.220  0.00411 ** 
## SpecimenSpecimen 1897 -0.110428   0.063000  -1.753  0.09422 .  
## SpecimenSpecimen 2005 -0.193912   0.068488  -2.831  0.01000 *  
## SpecimenSpecimen 2132 -0.096956   0.063000  -1.539  0.13874    
## SpecimenSpecimen 2134 -0.035943   0.063000  -0.571  0.57437    
## SpecimenSpecimen 2151 -0.163412   0.063000  -2.594  0.01694 *  
## SpecimenSpecimen 820  -0.088616   0.063000  -1.407  0.17417    
## lib_prepXT             0.005965   0.032208   0.185  0.85484    
## rxn_scale1X            0.060327   0.032208   1.873  0.07505 .  
## ---
## Signif. codes:  0 '***' 0.001 '**' 0.01 '*' 0.05 '.' 0.1 ' ' 1
## 
## Residual standard error: 0.08909 on 21 degrees of freedom
## Multiple R-squared:  0.4936, Adjusted R-squared:  0.2765 
## F-statistic: 2.274 on 9 and 21 DF,  p-value: 0.05814
```

### Beta diversity

Beta diversity (similarity between samples) was assessed by Bray-Curtis distance.

```
## 
## Call:
## adonis(formula = d ~ Specimen + lib_prep + rxn_scale, data = s) 
## 
## Permutation: free
## Number of permutations: 999
## 
## Terms added sequentially (first to last)
## 
##           Df SumsOfSqs MeanSqs F.Model      R2 Pr(>F)    
## Specimen   7   2.51299 0.35900 241.664 0.98140  0.001 ***
## lib_prep   1   0.01368 0.01368   9.207 0.00534  0.001 ***
## rxn_scale  1   0.00276 0.00276   1.859 0.00108  0.160    
## Residuals 21   0.03120 0.00149         0.01218           
## Total     30   2.56062                 1.00000           
## ---
## Signif. codes:  0 '***' 0.001 '**' 0.01 '*' 0.05 '.' 0.1 ' ' 1
```
